# Projection Neurons from Infralimbic (IL) to the Basolateral Amygdala (BLA) are Critical for the Expression of Social Familiarity induced Anxiolysis (SoFiA) in rats

**DOI:** 10.64898/2026.10.27.762180

**Authors:** Sreeparna Majumdar, Andrew R Burke, Erik T. Dustrude, Elizabeth A. Lungwitz, Katharine D. Andrews, Amy D Dietrich, William A. Truitt

## Abstract

Mental health is crucially linked to social behavior. Healthy social behavior involves learning to adapt emotional responses to social cues, for example learning to suppress anxiety through social familiarity, or social familiarity-induced anxiolysis (SoFiA). SoFiA is well documented and forms the basis for interpersonal therapy, however, the neural mechanisms of SoFiA are unclear. SoFiA is modeled in rats by employing the social interaction habituation (SI-hab) protocol.

Using SI-hab protocol, it has been determined that SoFiA represents social safety learning, which requires both anxiogenic stimulus and social familiarity during training sessions (5-6 daily SI sessions), and SoFiA expression is dependent on infralimbic subdivision (IL) of the ventromedial prefrontal cortex and the basolateral amygdala (BLA). Here we investigated the role of IL in SoFiA acquisition and expression, using chemogenetics to inhibit IL neurons. Male Wistar rats received bilateral viral injections targeting the IL or BLA and left undisturbed for 5-7 weeks to allow for viral expression. We chemogenetically inhibited via low doses of ip. clozapine-N-oxide (CNO) either the IL, the BLA, or the projections neurons from the IL to the BLA during expression of SoFiA in rats. We found that projection neurons from IL to the BLA are critical for the recovery SI behavior from the attenuation caused by the bright light (expression of SoFiA). Furthermore, CNO inhibited IL neurons with projections to the BLA and there were no off target effects of CNO on the behavioral measures. These results suggest an essential role of the IL-BLA pathway in social safety learning.

**Significance Statement:** The proposed research is relevant to public health because it will increase our systematic/mechanistic understanding of how social support improves the ability to manage anxiety responses, which will provide novel insights into our understanding of cognitive control of emotional behaviors. Impairments in social behaviors are the most prominent and disabling features of mental illness. This research will lead to increased understanding of the neural regulation of behavior and provide insights into ways to eventually reduce the burden of mental illness.

## Introduction

Anxiety disorder is one of the most common of psychiatric diseases with a 33.7% lifetime prevalence according to large population-based surveys (Bandelow and Michaelis, 2015) and it contributes to the etiology of major depression and substance abuse (Ressler and Mayberg, 2007; Koob, 2009; Tye et al., 2011). Treatment of anxiety disorder is mostly focused on a relatively non-specific treatment regime and includes psychotherapy or anxiolytic medications. Despite being valuable anxiety treatment options, psychotherapies have relatively high rate of attrition (e.g., Craske et al., 2019).

Social support can be anxiolytic and its quality depends on factors such as source of social support, frequency and context of social experience. Social sources comprising of great familiarity represent high potential for anxiolysis and social sources of low familiarity represents low potential for anxiolysis, and social isolation represents little or no potential for anxiolysis (Hawkley and Cacioppo, 2010; Dour et al., 2014). The highest anxiolytic quality of social support is achieved from interpersonal or group psychotherapies, where the objective is directed towards obtaining safety learning (Vervliet et al., 2013; Jacoby and Abramowitz, 2016). Clients with high levels of perceived social support have more positive outcomes of psychotherapy compared to low perception of social support (Ozbay et al., 2007; Umberson and Montez, 2010). The best predictor of success of these therapies depend on the therapeutic alliance and this social affiliation between the client and the therapist. (Fluckiger et al., 2018; Wu et al., 2024). In spite of unequivocal knowledge that strong social support corresponds with overall wellbeing and better health, and studies reporting involvement of prefrontal cortices in response to social support mediated attenuation of response to threat (Master et al., 2009; Coan et al., 2017; Berg et al., 2025), we still do not understand the underlying mechanisms clearly.

In the current study, we utilized an animal model that allows for manipulation of brain regions thought to underlie social support mediated reduction in perceived threat (anxiety-like) behavior in rats. While anxiety can be assessed through self-reports in human subjects, this method is inefficacious in rodents. Instead we expose rats to a bright light to capitalize on their innate aversion to brightly lit open spaces (File, 1980; Sajdyk and Shekhar, 1997). While there are different paradigms that measure potential threat behavior in animals; here, we focus on the rats native aversion towards social approach during bright light challenge in an open-field testing arena. The social interaction (SI) test is ideal for assessing potential threat behavior as it is a validated rat anxiety-related test that can be administered repeatedly and is not confounded by presence of a conspecific (File and Hyde, 1978; Sanders and Shekhar, 1995a, b). The SI test’s output measure, SI time, is determined as the amount of time the experimental rat spends initiating non-aggressive contact with the conspecific partner. SI time is inversely related to anxiety-like behavior.

A crucial aspect of healthy social behavior involves learning to adapt emotional responses to social cues, for example learning to suppress anxiety through social familiarity, or social familiarity-induced anxiolysis (SoFiA). SoFiA is modeled in rats by employing a social interaction habituation (SI-hab) protocol (Truitt et al., 2007; Lungwitz et al., 2014). Using SI-hab it has been determined that SoFiA represents social safety learning, which requires both anxiogenic/potential threat stimulus and social familiarity during SI training sessions. Previous work demonstrates that SoFiA expression is dependent on the infralimbic division (IL) of the ventromedial prefrontal cortex (vmPFC) (Lungwitz et al., 2014). Here the role of the IL, BLA and their circuitry in regulation of SoFiA expression were investigated using chemogenetic inhibition of the IL and/or the BLA during the SI-hab protocol. Inhibiting the IL, the BLA, and the IL efferents to the BLA disrupted expression of SoFiA-like behavior in rats, strongly establishing the involvement of IL-BLA pathway in the expression of SoFiA.

## Methods

### Animals

The subjects were male Wistar rats (Envigo Laboratories, Indianapolis, IN). For studies utilizing Designer Receptors Exclusively Activated by Designer Drugs (DREADDs), rats arrived at 3 weeks old weighing 30-75 grams. Rats arrived weighing 275-300 grams for all other experiments. Rats were individually housed in a temperature-controlled room (23°C) with 12:12 hour light-dark cycle with lights on at 0700h and *ad libitum* access to food and water. This study was carried out in accordance with the recommendations in the Guide for the Care and Use of Laboratory Animals of the National Institutes of Health. The Institutional Animal Care and Use Committee of the Indiana University Purdue University at Indianapolis approved all procedures.

### Social interaction (SI) test

Rats were acclimated to dim light, experimenter handling, and the behavioral testing apparatus before all behavioral experiments. The SI arena is a black Plexiglas square box (91.4 x 91.4 cm) with open top (30.5 cm high walls). Test and partner rats were acclimated to the dimly lit behavior room in their home cages for at least 45 mins prior to SI testing. During the 5 minute SI testing session, the test rat was placed in the open field box along with a conspecific partner that was age-, weight- and sex-matched. Each session was video recorded with an overhead camera. Social interaction (SI) time is the amount of time the test rat spent interacting with the familiar rat excluding aggression and sexual behavior. Seconds engaged in SI were quantified with ODLog (Macropod Software). Partner rats were used for maximum of two sessions in a day with at least 30 minutes gap between sessions. All SI testing was performed between 0900 and 1300 h.

Social interaction habituation (SI-hab) protocol comprises of multiple SI sessions repeated daily for at least five consecutive days. The SI test was performed with a social partner, either novel or familiar, under either non-threatening (dim light; 9-16 lux) or potential threat [bright light challenge (BLC); 500 (darkest corner)-1500 (center) lux] conditions. Lighting conditions (dim light or BLC), partner familiarity (novel or familiar partner) and pharmacological interventions are the variables that were controlled during SI-hab testing.

During SI testing, BLC was employed to induce perceived threat behavior, which consists of sudden change from dim red light (∼9 lux) to BLC (∼1000 lux, measured on the floor of the testing arena that is approximately 8.5 feet from the light source).

### Histology

Green fluorescent protein (GFP) or mCherry were the visual reporters of viral expression for control and DREADD viruses, respectively. Rats were perfused transcardially with saline and paraformaldehyde. Brains were removed, fixed in 4% paraformaldehyde (pH 7.34) for 90 min, washed in phosphate buffered saline (PBS), and then soaked in 30% sucrose until the brain sinks to the bottom. The brains were sliced coronally at 30 microns with every sixth section of the brain collected in a single well and stored in cryoprotectant (containing sucrose and ethylene glycol) at -20°C. For microinjections, the sections were stained with neutral red to aid in neuroanatomical landmark identifications

To visualize mCherry expression, sections were incubated with antibody, which specifically recognized mCherry (Abcam, ab 167453; 1:40000), overnight at room temperature. The next day, sections were washed with PBS, and then incubated with anti-rabbit secondary antibody (Vector Laboratories, 1:500) for 1 hour at room temperature. An ABC kit was used following PBS washes. Then brain sections were incubated in DAB (3’3’-diaminobenzidine) immunohistochemistry substrate (1:1000) for an hour. This substrate contained two components one being the DAB immunohistochemistry concentrate (reagent A) and the other was DAB immunohistochemistry reaction buffer (reagent B). Sections were washed in PBS and then DAB was used for 10 min. Chromagen reaction was quenched using 0.1M PB. Following washes, sections were mounted using 0.3% gelatin. All washes were done using 0.1M PBS.

To estimate the extent of viral spread or location of the microinjector tip, the brain atlas (Paxinos and Watson, 2007) and microscopes were used.

### Site-specific intracranial injections surgery

Rats were anesthetized with 3% isoflurane in 100% O_2_ with a flow rate of 0.6 L/min. A feedback-controlled heating pad was used to maintain the body temperature at 37°C. The head of the animal was placed in a stereotaxis frame and pretreated with lidocaine. For IL and BLA injections, following incision of scalp, a craniotomy was done and then dura was reflected, and pipettes/internal cannula were advanced to the desired stereotaxic coordinates of the target.

Bupivacaine (∼5 mg/kg) was administered intraincisionally during surgery for the BLA DREADD experiment. Following surgery, either buprenorphine or carpofen was injected subcutaneously for pain relief during recovery.

### Adenoviral injections surgery

Virus injections were performed using Hamilton syringe that were connected to a UltraMicroPump 3 driven by a Micro 4 MicroSyringe Pump Controller (World Precision Instruments). For anterograde expression of hM4D(Gi) recombinant adeno-associated virus AAV5-CAMKIIa-hM4D(Gi)-mCherry (Titer: 4.4x10^12 gc/ml) was injected (200 nl) at a rate of 80 nL/min. The virus was purchased from Addgene (Addgene 50477). CAMKIIa driven EGFP expression control AAV5-CAMKIIa-EGFP (Addgene 50469; Titer: 4.3x10^12 gc/ml) was used. CRE-recombinase expressing canine adenovirus-2 (CAV2; Titer: 1.1x10^12 gc/ml) with adeno-associated virus [pAAV5-hsyn-DIO-hM4D(Gi)-mCherry; Titer: 1.2x10^13 gc/ml] were injected in amygdala and IL, bilaterally, respectively. Cre-dependent Gi-DREADDs was obtained from Addgene (Addgene 44362) and CAV Cre was obtained from Plateforme de Vectorologie de Montpellier (PVM) Istitut de genetique moleculaire de montpellier (IGMM; Montpellier, France). Stereotaxis coordinates (in mm relative to bregma) for IL injections were AP (+3.2), ML (±0.5), DV (-5.0). For BLA injections, the coordinates were AP (-2.30), ML (+4.80), DV (-7.4).

### Intraperitoneal CNO injections

Clozapine-N-Oxide (CNO) was obtained from Tocris and diluted to the appropriate dose with 0.9% physiological saline, which was reconstituted prior to any injection. Various doses of CNO ranging from 0.25mg/kg to 3mg/kg injections were administered, intraperitoneally, 30 min prior to SI testing.

### Intracranial CNO injections

In order to attain quick, reversible, specific, selective and localized control, we pharmacologically intervened using designer receptor hM4D (modified version of Gi-coupled human muscarinic receptor 4) along with its exogenous ligand CNO to inhibit activity of neurons. Cannulae were implanted in BLA of rats previously injected with AAV5-CAMKIIa-hM4D(Gi)-mCherry in their IL. For BLA injections, following were the coordinates (in mm relative to bregma): AP (-2.30), ML (+4.80), DV (-7.4) at a 0° angle off the vertical plane. CNO (3μM, 100nL) was microinjected through the cannula 5 min prior to behavior testing.

### Patch clamp electrophysiology

Anesthetized male Wistar rats (∼250gms) were transcardially perfused with 25-30mL artificial cerebrospinal fluid (aCSF) having 93 N-methyl-D-glucamine (NMDG) among other usual components. Following decapitation, brains were dissected and coronal slices (350μm) were sectioned on Campden 7000smz-2 vibratomes (Lafayette Instrument Co, Lafayette, IN). The osmolarity of aCSF solution was adjusted to ∼315mOsm. Whole-cell current clamp recordings, using borosilicate glass electrodes (resistance 3-6mΩ), were performed at 30° C acute brain slices, which were visualized under Scientifica SliceScope Pro 6000 (Scientifica, Uckfield, UK) upright microscopes. Infralimbic cells labeled with mCherry were patched (n=4). 100pA current injections were applied. Recordings were done using aCSF and CNO (500nM). Action potential and membrane potential were recorded.

### Experimental design and statistical tests

To find the optimal dose of CNO that is absent of any off target effect on SI time, we first gathered baseline SI times in dim light in the absence of CNO, then we administered the 5 doses of CNO repeatedly one time per day to the same rats (n=6) in a random order (Figure 1). These data were analyzed with a repeated measures 1-way ANOVA and followed up by Dunnett’s (BLC comparison) and Fisher’s least squared differences (doses comparisons) post hoc tests.

**Figure 1:**
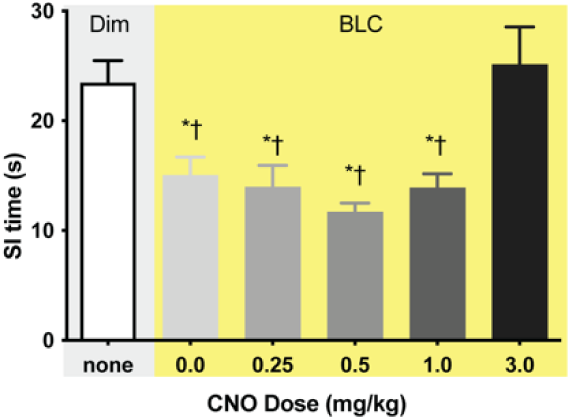
Rats exhibit BLC effect at lower systemic CNO doses with no off target effects. *indicates significantly different from baseline (Dunnett’s p ≤ 0.0045). † indicates significantly different from 3mg, LSD p ≤ 0.0015. The light grey block indicates dim light condition while the yellow block indicates anxiogenic stimulus (BLC). X axis represents different doses of CNO between 0 and 3 mg/kg BW of animals. Y axis represents SI (in seconds) time ± SEM (n=6). Abbreviations: clozapine-n-oxide (CNO), bright light challenge (BLC).

Rats (n=5) pretreated with Gi-DREADD virus underwent eleven days of SI-hab protocol (ST1-ST11) with CNO administered on social training day 9 to determine the role of the vmPFC (IL) on SoFiA (Figure 2). These data were analyzed with a repeated measures 1-way ANOVA followed up by Tukey’s (day comparisons) and Fisher’s (drug comparisons) post hoc tests. A separate group of rats (n=7) were pretreated with the Gi-DREADD virus and were administered ip. CNO or saline 30 mins prior to SI testing under dim light with a novel partner (baseline conditions). Of these rats, only three rats were tested with both saline and CNO on separate days, yielding n=5/group. Two were tested with saline and then CNO, whereas one was tested with CNO and then saline. These data were analyzed with a Student’s t-test.

**Figure 2:**
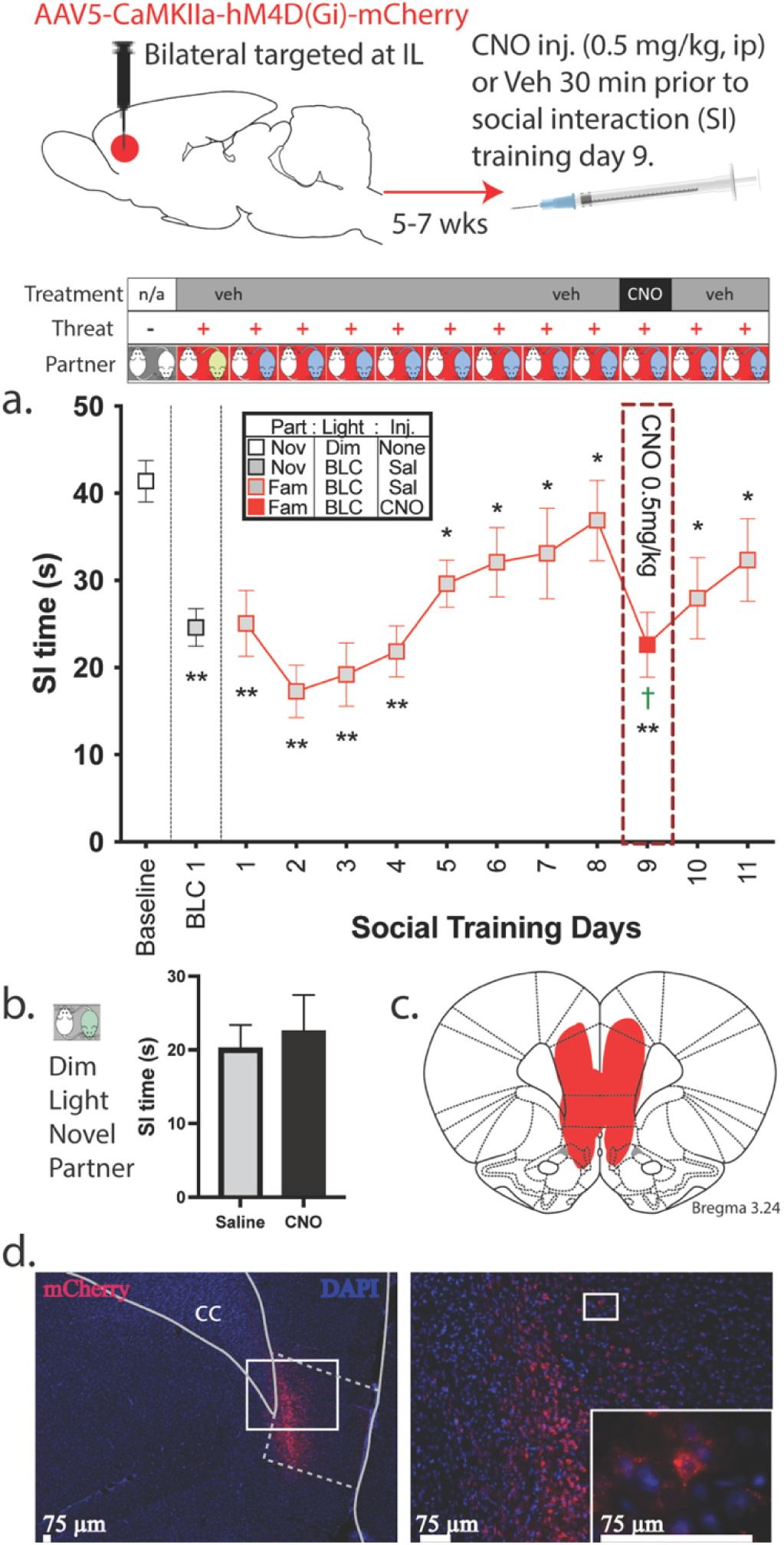
CNO disrupts SoFiA expression in rats with hM4D-DREADDs in vmPFC. Top, procedural schematic of experimental design. A, SoFiA acquisition and expression, which is blocked by chemogenetic inhibition of the IL via CNO. * indicates significantly different from ST2, Tukey’s p ≤ 0.03. ** indicates significantly different from baseline Tukey’s p ≤ 0.0354. † indicates CNO significantly different from ST8, 11 Fisher’s LSD p ≤ 0.014. X axis represents social training days, Y axis represents SI (in seconds) time± SEM (n=5). B, SI time during baseline conditions in separate group of rats n=5/group. C, extent of the spread of mCherry expression from the viral injection site targeted at the IL. D, representative image of an injection site in IL of rat injected with CAMKII-GiD. The region of interest is in dashed outline. Right panel is image at higher magnification from within white rectangle found in left panel. Lower right image from rectangle in right panel shows double labeled cells with DAPI and mCherry. Abbreviations: infralimbic cortex (IL), clozapine-n-oxide (CNO), partner (Part), novel (Nov), familiar (Fam), bright light challenge (BLC), corpus collosum (cc), 4′,6-diamidino-2-phenylindole (DAPI).

To determine the extent to which the BLA is required for SoFiA expression, we pre-treated the BLA with CaMKIIa-Gi or a control virus in a separate group of rats (n=6/group; Figure 3). After investigating the BLC with a 2-way ANOVA followed by Bonferroni post hoc tests, we utilized a repeated measures 1-way ANOVA on each treatment group separately because were interested in how CNO impacts SoFiA behavior after it has been expressed. One-way ANOVAs were followed up by Dunnett’s (day comparisons) and Fisher’s (drug comparisons). To ensure that BLA inhibition did not alter baseline SI time, we counterbalanced CNO and vehicle treatment order of administration at the end of the experiment and compared SI times with a t-test.

**Figure 3:**
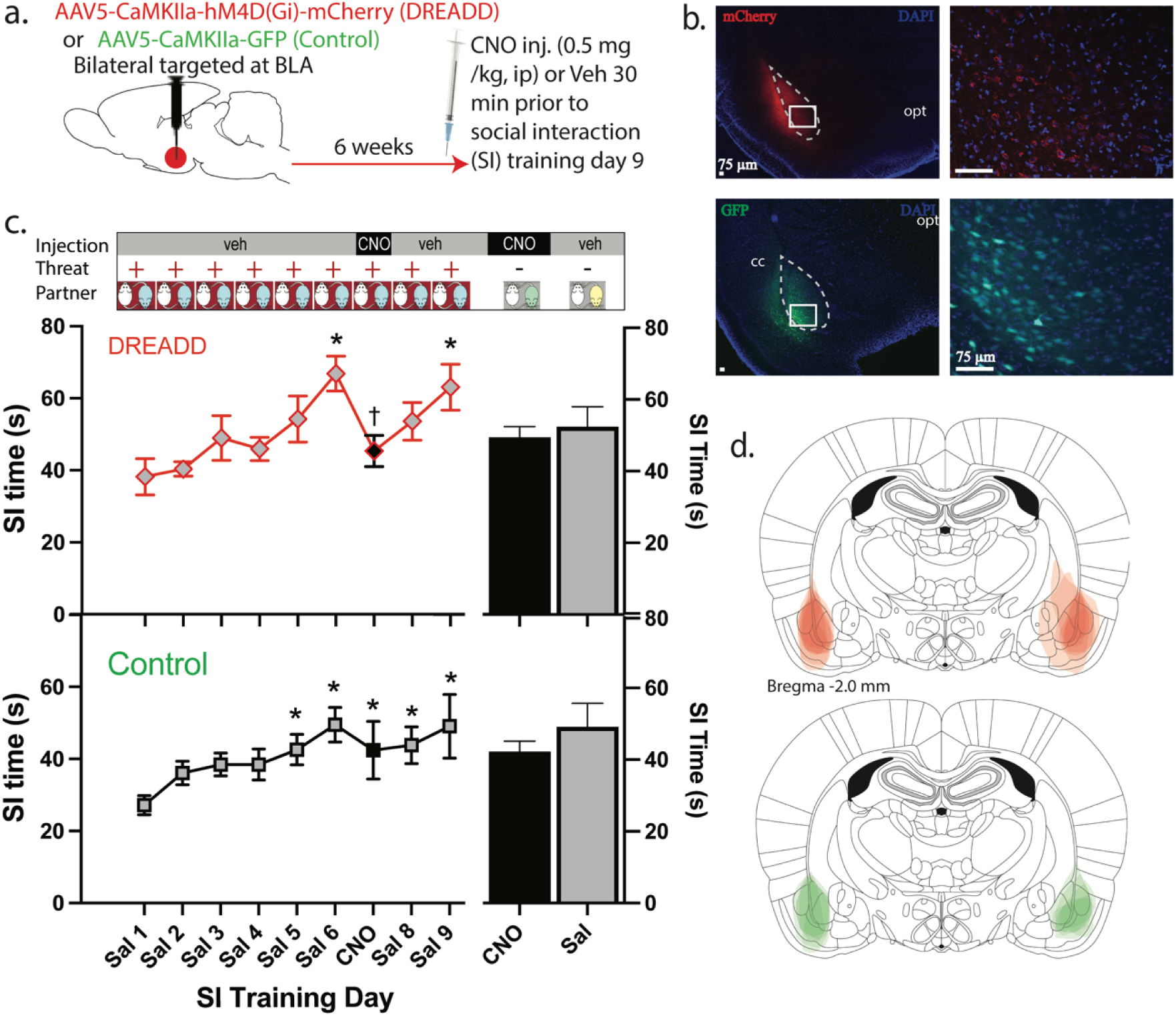
CNO disrupts SoFiA expression in rats with hM4D-DREADD in BLA. A, procedural schematic of experimental design. B, representative image of BLA of rat injected with CAMKII-GiD. The region of interest is in dashed outline. Right panel is image at higher magnification from within white rectangle found in left panel. Lower right image from rectangle in right panel shows double labeled cells with DAPI and mCherry. C, SoFiA acquisition and expression, which is blocked by CNO. *indicates significantly different from ST2, Dunnets p ≤ 0.01. †indicates CNO significantly different from Sal 6 (ST6), Fisher’s LSD p < 0.001. X axis represents social training days, Y axis represents SI (in seconds) time± SEM (n=6/group). D, extent of the spread of mCherry (red) and GFP (green) expression from the viral injection site targeted for each rat. Abbreviations: basolateral amygdala (BLA), clozapine-n-oxide (CNO), anxiety stimulus (anx stim), corpus collosum (cc), optic fiber (opt), 4′,6-diamidino-2-phenylindole (DAPI).

To identify the pivotal targets of vmPFC (IL) projection neurons are critical for SoFiA we injected two different types of viruses in rat IL, with CaMKIIa-Gi in one group (n=7) and the control virus in the other group (n=5; Figure 4). Following SoFiA acquisition CNO was delivered intracranially into the BLA, through the implanted guide cannula 5 minutes prior to social training day 7. These data were analyzed with a repeated measure 2-way ANOVA followed up by Tukey’s (BLC comparisons) and Dunnett’s (day comparisons) post hoc tests. The percent inhibition of SoFiA expression (Figure 5f) was analyzed with a repeated measures 2-way ANOVA followed up by Tukey’s for post hoc comparisons.

**Figure 4:**
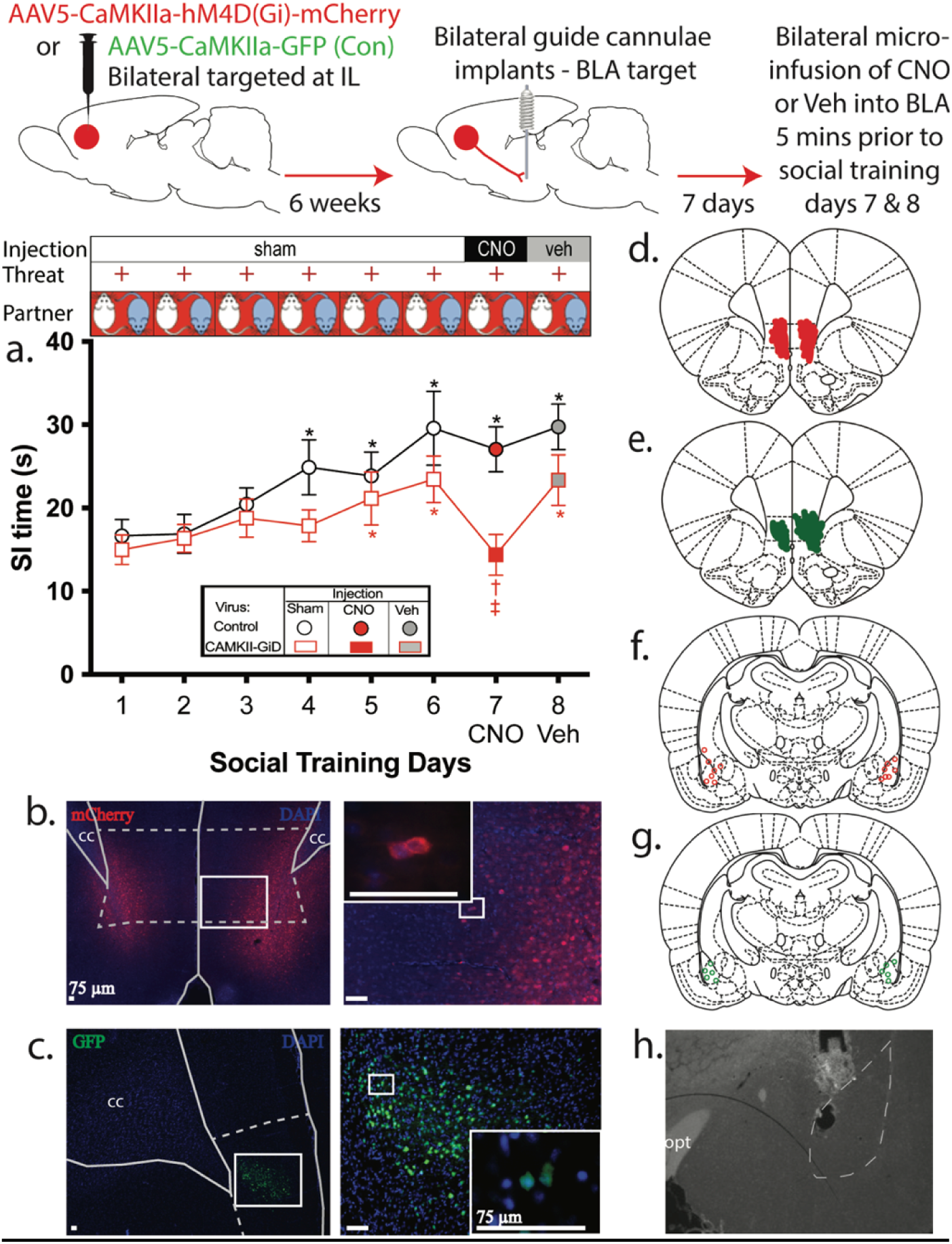
Intra-BLA CNO blocks SoFiA expression in rats with vmPFC hM4D-DREADDS. Top, procedural schematic of experimental design. A, SoFiA acquisition and expression for rats pretreated with the DREADD virus (CAMKII-GiD) virus or control virus (n=5-7/group) in the IL and fixed with guide cannualae aimed at the BLA. *indicates significantly different from ST1, Dunnett’s p ≤ 0.021 †indicates significantly different from ST6 & 8, Tukey’s p ≤ 0.0007 ‡indicates significantly different from control, Fisher’s LSD p = 0.0013. B, representative image of an injection site in IL of rat injected with CAMKII-GiD. C, representative image of an injection site in IL of rat injected with control virus. For B & C, right panels are image at higher magnification from within white rectangle found in left panel. Corner images shows double labeled cells with DAPI and mCherry at higher magnification. The region of interest is in dashed outline. D, extent of the spread of mCherry expression from the viral injection site targeted at the IL for the CAMKII-GiD group. E, extent of the spread of GFP expression from the viral injection site targeted at the IL for the control virus group. F, Location of the CNO or veh microinjection sites in the BLA for the CAMKII-GiD group. G, Location of the CNO or veh microinjection sites in the BLA for the CAMKII-GiD group. H, representative image showing injection site within the BLA. The region of interest is in dashed outline. Abbreviations: infralimbic cortex (IL), basolateral amygdala (BLA), clozapine-n-oxide (CNO), anxiety (Anx), Stimulus (Stim), bright light challenge (BLC), corpus collosum (cc), 4′,6-diamidino-2-phenylindole (DAPI), optic tract (opt).

**Figure 5:**
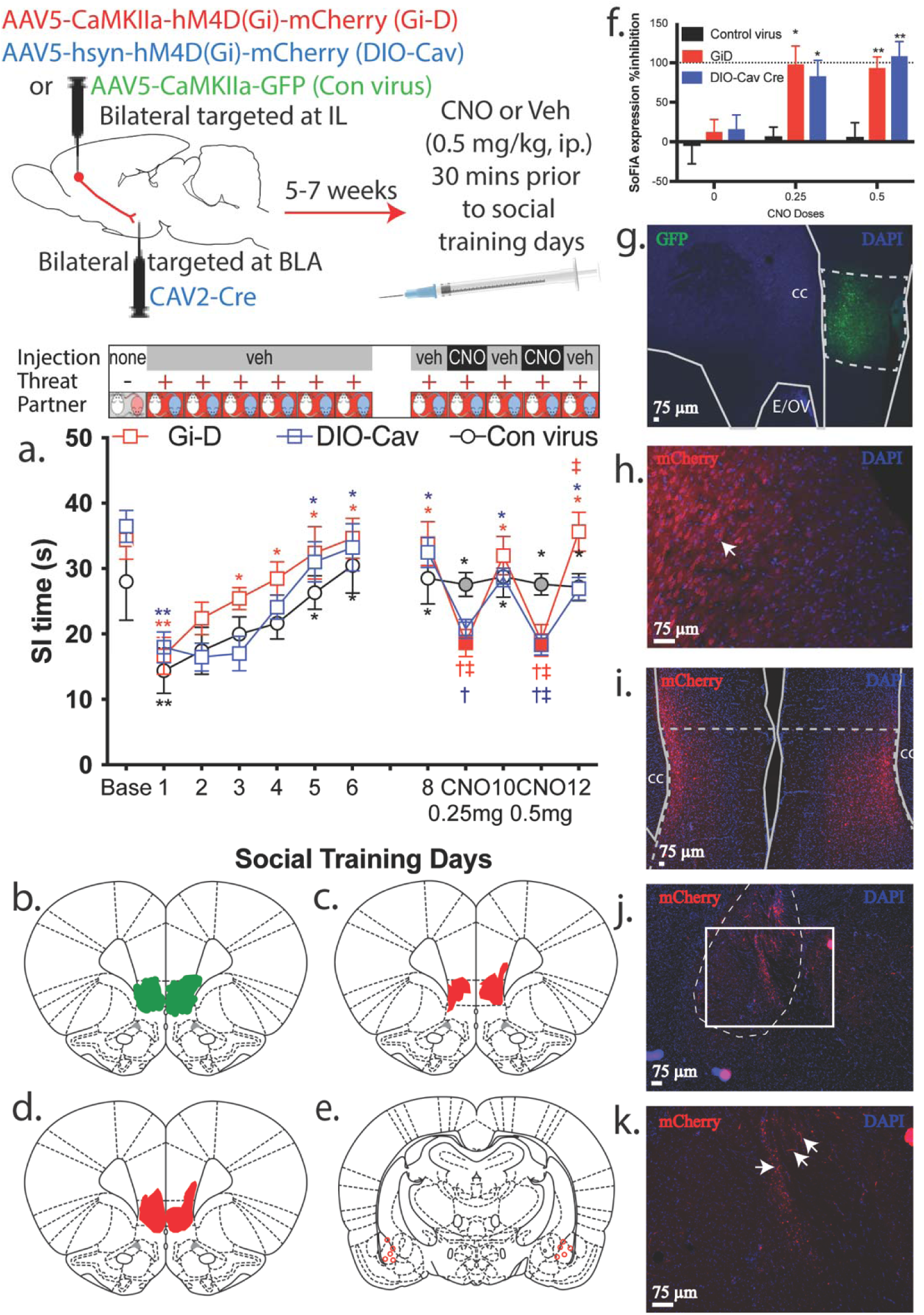
CNO inhibits SoFiA expression in rats with intersectional DREADD virus targeting IL projections to the BLA. Top, procedural schematic of experimental design. A, SoFiA acquisition and expression for rats pretreated in the IL with the DREADD virus (CAMKII-GiD) virus, intersectional DREADD virus (DIO-Cav) or control virus and then receiving systemic CNO and vehicle injections prior to each social training day (n=4-5/group). *indicates significantly different from ST1, Dunnett’s p ≤ 0.037. **indicates significantly different from baseline, Dunnett’s p ≤ 0.0001 †indicates significantly different from ST8, 10, 12, Tukey’s p ≤ 0.007 ‡indicates significantly different from control, Fisher’s LSD p ≤ 0.04. B, extent of the spread of GFP expression from the viral injection site targeted at the IL for the control virus group. C, extent of the spread of mCherry expression from the viral injection site targeted at the IL for the CAMKII-GiD group. D, extent of the spread of mCherry expression from the viral injection site targeted at the IL for the DIO-Cav group. E, site of pretreatment with Cav2-Cre virus in the BLA for the DIO-Cav group. F, Percent inhibition of SoFiA expression. Zero mg/kg CNO failed to alter the percent inhibition, whereas in both GiD and DIO-CAV Cre groups, the percent inhibition for both 0.25 mg CNO (Tukey’s p = 0.009 and p = 0.025 for Gi-D and Cav-Cre, respectively), and 0.5 mg CNO (Tukey’s p = 0.014 and p = 0.002 for Gi-D and Cav-Cre, respectively) doses were significantly different from control virus. G, representative image of an injection site in IL of rat injected with control virus. The region of interest is in dashed outline. H, representative image of an injection site in IL of rat injected with CAMKII-GiD with arrow pointing to double labeled cell body. I, representative image of an injection site in IL of rat injected with DIO virus. The region of interest is in dashed outline. J, representative image of CAV2-Cre injection site in the BLA. K, higher magnification of I from within white rectangle with arrows pointing to fibers stained with mCherry showing the virus traveled from the IL to the BLA. K, Representative image of injection site of CAV2-Cre within the BLA (note that the damage is light because tissue is collected 8 weeks post microinjection). The region of interest is in dashed outline. Abbreviations: infralimbic cortex (IL), basolateral amygdala (BLA), clozapine-n-oxide (CNO), Stimulus (Stim), bright light challenge (BLC), corpus collosum (cc), Ependymal layer/olfactory ventricle (E/OV), 4′,6-diamidino-2-phenylindole (DAPI), optic tract (opt).

To confirm whether IL projections to BLA regulate SoFiA expression we used intersectional chemogenetics, which targeted the IL projections to the BLA (Figure 5). There were three different groups (n=4-5/group) based on the type of virus Gi-D, DIO-Cav, and Con Virus. The acquisition of SoFiA data were analyzed with a repeated measures 2-way ANOVA followed up by Tukey’s (BLC comparisons) and Dunnett’s (day comparisons). The ability of CNO-induced inhibition of the IL and IL projections to the BLA was analyzed by a separate repeated measures 2-way ANOVA followed up by Fisher’s (virus comparisons) post hoc tests.

To verify function of the Cre-dependent DREADD virus, we conducted whole-cell voltage clamp recordings from mCherry+ IL cell bodies in hsyn-DIO-GiD-IL/CAV-Cre-BLA pre-treated rat brain slices under bath application of CNO (Figure 6). These comparisons were made by a Student’s t-test (n=3 cells).

**Figure 6:**
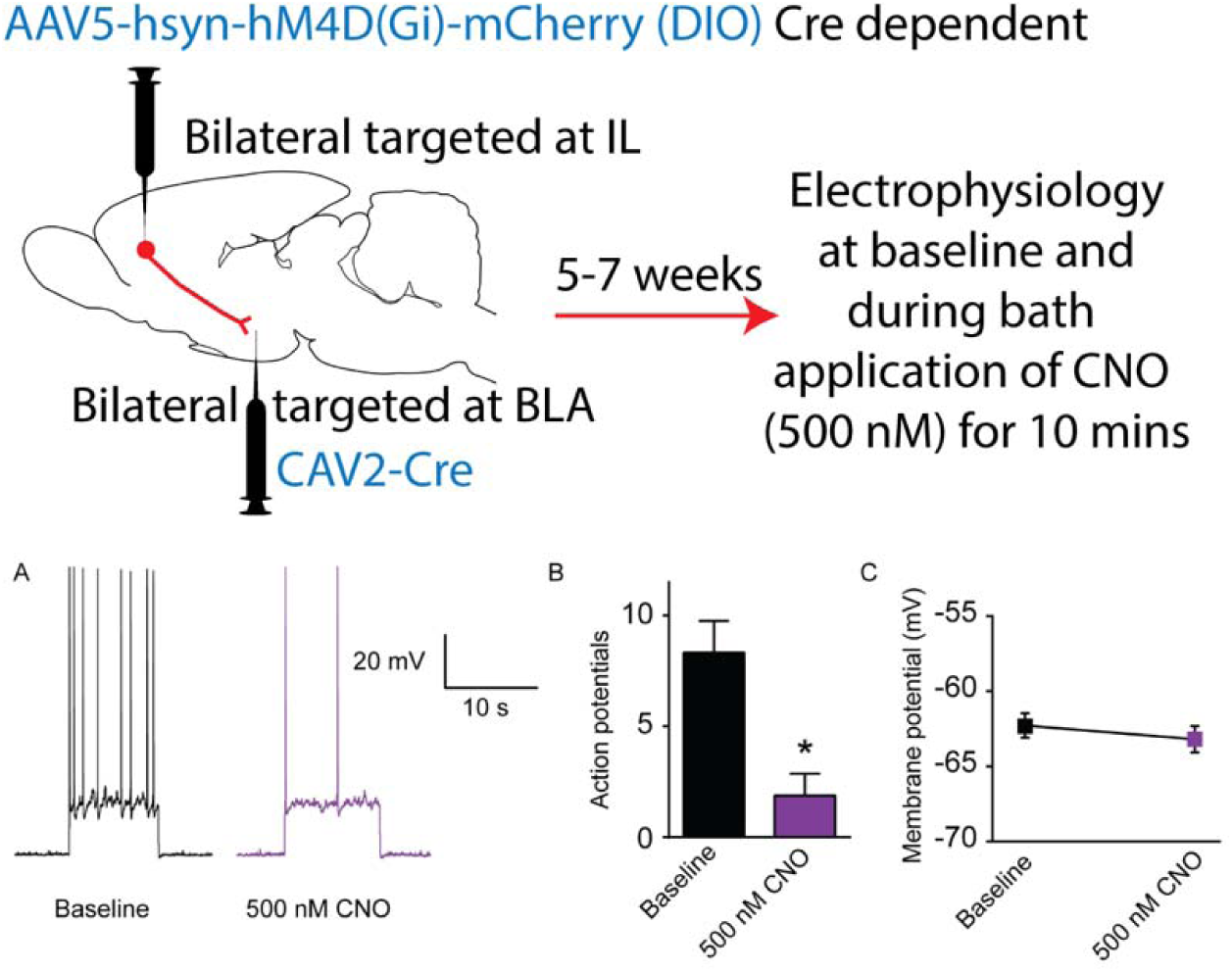
Functional connectivity between IL and BLA terminals was validated by whole-cell voltage clamp recording. Top, procedural schematic of experimental design. A&B, Whole-cell voltage clamp recording shows that bath application of CNO (500 nM) decreases the number of AP in hsyn-DIO-GiD+ rats when compared to baseline. A, Electrophysiological traces showing AP during baseline recording and CNO treatment. B, The graph shows that number of AP is significantly reduced with CNO (n=3 cells). * indicates Student’s t test, p = 0.006. (C) The membrane potential is comparable during baseline and CNO treatment. Abbreviations: infralimbic cortex (IL), basolateral amygdala (BLA), clozapine-n-oxide (CNO).

All data were analyzed using GraphPad Prism 6.0 or 10 (La Jolla, CA) and all data are presented as mean ±SEM with alpha set at 0.05 to be considered statistically significant.

## Results

### Low doses of CNO are optimal for BLC SI-hab testing

Since CNO has been reported to have off-target effects, our goal was to identify an optimal CNO dose critical for our experiments on social interaction-anxiety challenge assay. In naïve control (no viral expression/ lacking viral expression) rats, we assessed SI time during 5 different doses of CNO (0-, 0.25-, 0.5-, 1-and 3 mg/kg ip. 30 mins prior to test) under the BLC (Figure 1). We found that lower doses of CNO (0, 0.25-, 0.5- and 1mg/kg) significantly decreased the SI time from baseline SI [repeated measures one way ANOVA (F _2,10_)=18.46, p = 0.0005, Dunnett’s p ≤ 0.0045] and not significantly different from each other. The higher CNO dose (3 mg/kg BW) blocked the BLC effect and allowed for significantly increased SI time than rest of the lower CNO doses (0 to 1.0 mg/kg) (Fisher’s LSD p ≤ 0.0015). This suggests that lower doses of CNO (0 to 1.0 mg/kg) did not have any off-target effects on SI behavior (anxiety-like behavior). The subsequent chemogenetic studies utilized 0.25 and 0.5 mg/kg.

### CNO disrupts SoFiA expression in rats with hM4D-DREADD in vmPFC

Rats were injected with inhibitory DREADD virus (CaMKIIa-hM4D-mCherry) targeting the IL division of the vmPFC (figure 2 top schematic). These rats were exposed to SI-hab (Figure 2a top). Overall, SI time was affected by BLC and social familiarity [one-way ANOVA, (F _3, 12_)=9.95, p = 0.0015]. Compared to baseline BLC significantly reduced SI time during BLC 1 and during the first 4 days of social training (Tukey’s p ≤ 0.0354). Additionally, social familiarity significantly increased subsequent SI times over the next 4 social training days compared to the initial social training days (Tukey’s p ≤ 0.03). Thus, rats were expressing SoFiA from ST 5 through ST 8. On ST 9 an injection of 0.5 mg/kg CNO produced a reduction in SI time compared to the prior ST day (Fisher’s LSD p =0.006), indicating a disruption of SoFiA. When tested without further injection of CNO on ST 10 and 11 (Fisher’s LSD p ≤ 0.014), the SI time was similar to previous ST days and no longer significantly different from baseline.

Furthermore, among rats pretreated with the DREADD virus, there was no difference between CNO and saline under baseline conditions (dim light and novel partner; t-test p > 0.05; Figure 2b). The maximum area of the viral expression is depicted in Figure 2c, with representative images shown in Figure 2d. These findings suggest that CNO administration in these rats blocks SoFiA without altering SI times under baseline conditions.

### CNO disrupts SoFiA expression in rats with hM4D-DREADDs in BLA

Rats were injected with CaMKIIa-hM4D(Gi)-mCherry (DREADD) or CaMKIIa-GFP (Control) targeting BLA (Figure 3a) resulting in robust labeling of mCherry and GFP neurons (both cell bodies and axons) in the BLA (Figure 3b). Following baseline SI, these Gi-Dreadds+ rats underwent nine days of the SI-hab protocol (Sal 1-Sal 9; Figure 3c top) and CNO was systemically administered on the 7^th^ day (CNO). Controls exhibited significant change over time (F_9,45_=4.385, p = 0.004) with days Sal 5, Sal 6, CNO, Sal 9, and Sal 9 greater that Sal 1 (Dunnett’s, p ≤ 0.023; Figure 3c bottom). The DREADD group also showed a significant change over time (F_9,45_ = 4.720, p = 0.002) with days Sal 6, and Sal 9 being significantly greater. For the DREADD group only, the CNO day resulted in a significant reduction in SI time compared to Sal 6 and Sal 9 (Fisher’s p < 0.001; Figure 3c top), whereas the control virus treated rats had no significant reduction during CNO administration. These results suggest that the BLA is critical for expression of SoFiA. We found no effect of CNO during baseline conditions (dim light & novel partner; Figure 3c right), suggesting that the effects of CNO on DREADD and Control rats are unique to BLC conditions. The BLC reduced SI time in when comparing baseline to ST1 for Controls (t=5.05, p = 0.004) and DREADD (t=2.77, p = 0.040) rats (data not shown).

### Intra-BLA CNO blocks SoFiA expression in rats with vmPFC hM4D-DREADDs

To identify the pivotal targets of vmPFC (IL) projection neurons that regulate (or are critical for) SoFiA expression, rats were exposed to two different types of viruses in the vmPFC (IL). Rats were injected with either CaMKIIa-hM4D (Gi) or a CaMKIIa-EGFP control virus 6 weeks before being implanted with bilateral guide cannulae aimed at the BLA (Figure 4 top). Both groups underwent SI-hab to acquire SoFiA under sham injection conditions (Figure 4a top).

After SoFiA was acquired in both groups, CNO was delivered intracranially, through the implanted guide cannulae into BLA 5 minutes before SI test on ST7. There was a significant main effect of ST days and a significant interaction in the repeated measures two-way ANOVA, (F_7,70_)=12.99, p < 0.0001 and (F_7,70_)=3.31, p = 0.004, respectively. (Figure 4a). Initially both groups displayed a significant decrease in SI time from baseline SI (Tukey’s p ≤ 0.0043) on ST1 and ST2 compared to baseline (data not shown). Both control and CAMKII-GiD showed increased ST times compared to ST1. Control exhibited significantly greater SI time on ST4 through 8 (Dunnett’s p ≤ 0.021) and CAMKII-GiD exhibited significantly greater SI times on ST6, 6 and 8 (Dunnett’s p ≤ 0.019), suggesting SoFiA expression. The viral treatment groups displayed differential response to CNO. In CAMKII-GiD rats SI times following CNO was significantly decreased compared to control (Fisher’s LSD p = 0.0013) and from ST6-SI time in the CamKII Gi-D rats, following intracranial injection into BLA (Tukey’s p = 0.0007). In the CamKII Gi-D rats, the SI time returned back to pre-inhibition levels the following day (ST8) after vehicle microinjection (Tukey’s p = 0.0006) in CamKII Gi-D rats. However, the control virus injected rats did not exhibit any differential response to either CNO or vehicle microinjections into BLA.

### vmPFC-BLA neurons involved in SoFiA expression

Intersectional chemogenetics was used to confirm IL projections to BLA regulate SoFiA expression. There were three different groups based on the type of virus injection into the IL: control virus (CAMKIIa-EGFP), Gi-D (Cre independent/ CAMKIIa-hM4D(Gi)-mCherry) and DIO-Cav (hsyn-DIO-hM4D(Gi)-mCherry and CAV2 Cre), the latter allowing expression of DREADDs (injected into IL) in Cre expressing cells (CAV2 Cre injected into BLA; Figure 5 schematic). Following the usual SI-hab protocol (Figure 5a top), we analyzed baseline and ST1-6 first (SoFiA Acquision). There was a main effect of social training days [2-way RM ANOVA (F _6, 60_) =35.92, p < 0.0001] and all three groups displayed BLC effect, where the SI time was significantly decreased from baseline SI (Dunnett’s p ≤ 0.0001) on ST1 (Figure 5a). After this exposure to familiar conspecific resulted in a significant increase in SI time (i.e., SoFiA acquisition) from ST1 on ST4-6 for Gi-D, ST5 and 6 for DIO-Cav, and ST5 and 6 for control virus rats (Dunnett’s p ≤ 0.0005). All three groups continue to express SoFiA until ST8, where there is no differential effect of social training days on SI time among groups (Dunnett’s p ≤ 0.0001).

Following acquisition of SoFiA, all three groups were challenged with two different doses of CNO either with lower CNO dose (0.25 mg/kg) on ST9 or with higher CNO dose (0.5 mg/kg) on ST11 in a counterbalanced design. The 2-way RM ANOVA for this part of the experiment (ST8-12) suggested a significant main effect of day [(F _4.44, 44.4)_ = 27.62, p < 0.0001] and a significant day by virus group interaction [(F _8.88, 44.4_) = 3.13, p < 0.0054]. Challenging with both doses of CNO significantly decreased the SI time from days prior and post CNO treatment (Fisher’s LSD p ≤ 0.0088). Additionally, the CNO treatment significantly reduced the SI time compared to the control group (Fisher’s LSD p ≤ 0.0305), which remained unaltered by the CNO injections. For the percent change in inhibition, there was a significant main effect of virus (F _2,10_ = 5.018, p = 0.0310) and a significant virus by drug interaction and (F_4, 20_ = 3.624, p = 0.0223) respectively (Figure 5f). In both Gi-D and DIO-Cav groups, the percent inhibition for both 0.25 mg CNO (Tukey’s p ≤ 0.0098 and p ≤ 0.0246 for Gi-D and Cav-Cre, respectively), and 0.5 mg CNO (Tukey’s p ≤ 0.0139 and p ≤ 0.0023 for Gi-D and Cav-Cre, respectively) doses were significantly different from control virus.

These results suggest that inhibiting neurons in the IL or a subset of the IL neurons that project to the BLA both result in disruption of SoFiA expression. Also, the level of inhibition mediated through targeting selective IL projections to the BLA appears to equal that of inhibition level of pan IL neurons.

### Electrophysiology analysis to establish functional connectivity between IL and BLA

We sought to verify function of Cre-dependent Gi-DREADD virus by utilizing whole-cell voltage clamp recordings from IL cell bodies from rats that were pre-treated 5-7 weeks earlier with hsyn-DIO-hM4D(Gi)-mCherry into the IL and CAV2-Cre into the BLA. Ten minutes’ perfusion of 500 nM CNO significantly reduced the number of action potentials generated by a 10 s current injection of 100 pA (t-test, p = 0.0055, t = 7.203, df = 3) (Figure 6B). Resting membrane potential was unaffected by CNO application (t-test, p = 0.3948, t = 0.9909, df = 3) (Figure 6B). These data indicate functional connectivity between IL cell bodies and terminal fields within the BLA, and that activity of Gi-DREADD expressing IL cells can be reduced by CNO treatment. The membrane potential during baseline and CNO treatment remains comparable (Figure 6C)

## Discussion

This investigation of the impact of chemogenetic inhibition on the vmPFC and the BLA indicated that these regions are essential for the expression of a recovered SI time caused by a familiar social partner during the SI-hab experiments, which is a model of SoFiA expression (Truitt et al., 2007; Lungwitz et al., 2014; Majumdar et al., 2018; Race et al., 2021; Burke et al., 2025). While screening for off target effects of the CNO, we discovered that 3 mg/kg of CNO blocks the BLC suppression of SI time, but lower doses (e.g. 0.5 mg/kg) had no effect on SI time. During SI-hab, we observed a pronounced reduction in SoFiA expression following chemogenetic inhibition of either IL or BLA, indicating the critical involvement of these brain regions in the recovery of SI time over SI-hab sessions. It is important to note that CNO did not affect SoFiA expressions in rats injected with control virus further confirming a lack of off-target CNO effects in the doses used here. Furthermore, our findings demonstrate that the inhibition of IL projections to the BLA through intra BLA infusion of CNO or via intersectional chemogenetics significantly blocked the increase in SI time observed during SI-hab, underscoring the interconnected nature of these regions in regulating the expression of SoFiA. Our in vitro experiments suggest an inhibitory action of intersectional chemogenetics on IL cells. Collectively, these results suggest that the expression SoFiA in this SI-hab rat model requires intact IL neurons that project to the BLA.

### CNO effects were specific to DREADDs

Considering the off target effects of CNO and its metabolites (reviewed in Rodd et al., 2022), we investigated whether systemic CNO would impact our dependent variable, SI time, prior to conducting any DREADD experiments. Our results showed that only at the highest dose tested, 3 mg/kg, did CNO increase SI time under the bright light. Social interaction time in a brightly lit area has been found to increase during the systemic administration of other anxiolytic drugs (Depoortère et al., 2019). When previously testing lower doses of clozapine in this brightly lit SI test, there was no effect on social interaction time (Millan et al., 1999). Further support for the lack of off-target effects can be found in the control virus groups in Figure 3. Also, the virus expression in the BLA did not impact SI time during baseline conditions with a dim light and novel partner (Figure 2b & 3c bar graphs). Thus, we assume that the differences in SI time are a result of the IL or BLA inhibition, rather than from off-target CNO effects.

### vmPFC and BLA in SoFiA

Our past work found that local pharmacological inhibition of vmPFC (IL) (Lungwitz et al., 2014) or disruption of local inhibitory network of the BLA (Truitt et al., 2007) disrupted SoFiA expression and acquisition, respectfully. The current study utilizing DREADD technology suggest the CAMK2 neurons of vmPFC or the BLA are necessary for SoFiA expression. While spread of virus to areas adjacent to these structures cannot be completely ruled out, our findings are in line with other reports showing that the frontolimbic circuit plays an essential role in social behavior and emotional processing (Adolphs, 2003; Phelps and LeDoux, 2005; Felix-Ortiz et al., 2016). It appears that CAMK2 neurons are selectively involved in SoFiA expression as inhibition of these neurons did not alter social interactions in rats that had not learned SoFiA. Additionally, inhibition of these neurons did not alter acute responses to the bright light challenge further suggesting their roles are more likely involved with enforcing the learned SoFiA behavior.

Considering the IL has strong projection to the BLA (Bloodgood et al., 2018; Huang et al., 2020) via CAMK2 neurons, we also determined that IL projections to the BLA are equally necessary for expression of SoFiA. Using intersectional chemogenetics, we demonstrate that targeting a subset of IL CAMK2 neurons that project to the BLA was as effective at blocking SoFiA expression as targeting the population of IL CAMK2 neurons, where CNO was effective at reducing SoFiA expression in rats with either IL CAMK2 DREADD expression or DREADD expression only targeted by intersectional chemogenetics. While the intersectional chemogenetics group (DIO-CAV) was designed to target only IL neurons that project to the BLA, it is possible that some spread of virus may have reached IL efferents from adjacent areas or via IL efferents passing through the area of the BLA. In addition to BLA (Bloodgood et al., 2018) IL projects to other amydalar regions such as CeA, intercalated nucleus of amygdala (McDonald et al., 1996), and basal medial amygdala (BMA) and IL efferents to the bed nucleus of the stria terminalis (BNST) and hypothalamus (Vertes, 2004). It is possible that any of these IL-efferents could also play a role on reducing SoFiA expression. Murine studies have shown that IL cells project to BMA (Adhikari et al., 2015) and may be important for regulating anxiety like responses. Therefore, this information could be taken into consideration while interpreting the present data. However, when we targeted IL efferents in the BLA by targeted injections of CNO into the BLA, we also observed robust reduction of SoFiA expression, suggesting that these IL-projections to more distal structures may not be as crucial to the SoFiA expression as the IL projections to the BLA.

The connection from the IL to the BLA was confirmed in the current study using intersectional chemogenetics and in vitro electrophysiology during bath application of CNO (Figure 5 and 6).

### Overall Conclusions and Future Directions

Taken together, these results collectively provide evidence to support that IL, BLA and IL efferents to the BLA are critical for SoFiA expression. It is not clearly understood whether regulation of this specific IL-BLA pathway would be sufficient for SoFiA acquisition, social memory, or the reduction in SI time caused by the BLC. These results, however, are limited to males mostly because of the observation that SI time is attenuated by estradiol in ovariectomized females (Koss et al., 2004). Considering the importance of using both sexes in basic research (e.g., Shansky, 2019), moving forward females will be tested in the SI-hab model of SoFiA or the model will be adapted to include both sexes.

The mechanism by which bright light reduces SI time remains elusive and it was interesting, if not unexpected, that inhibiting the BLA did not suppress the anxiety-like behavior induced by BLC. Our lab has also confirmed that muscimol injections into the BLA is ineffective in reducing anxiety-like behavior during BLC (Burke, et al., 2024). It is possible that IL receives potential threat information from BNST, which perceives the anxiety signal via subregions of amygdala (e.g. CeA). It is probably the pBNST that is involved in perceived threat response to the bright light (Dong et al., 2001; Walker et al., 2009; Yamamoto et al., 2018). As the BNST receives projections from BLA (Dong et al., 2001), a circuit between these regions may underlie the inhibition of social interaction by bright light. However, the BLA by itself has been implicated in regulation of anxiety and emotional memory (Davis, 1994; McGaugh et al., 2002) and inhibiting and activating BLA decreases and increases anxiety-like behavior. Future studies investigating the role of these regions in perceived threat caused by bright lights are needed.

Another possibility is that IL-BLA projections activate a prosocial output of the BLA that is activated during SoFiA expression, and blocking either BLA output neurons directly or the IL-activation of this BLA prosocial output disrupts the SI increase observed during SoFiA expression.

There is some research suggesting that the IL projection to the BLA impairs social interaction behavior in a three-chamber social assay where mice interact with a confined social partner (Huang et al., 2020). Furthermore, the BLA projections to the IL (Felix-Ortiz et al., 2016), and the BLA alone (Garrido Zinn et al., 2016) are required for recall of social memory. Thus, it is also possible that by manipulating IL, BLA, or the IL projections to the BLA, we are interfering with the rat’s recall of the familiar partner rat. It will be prudent to investigate the role of these regions in social memory in the future as well as investigating SoFiA in brain regions involved social memory, such as the ventral hippocampus (Felix-Ortiz and Tye, 2014; Okuyama et al., 2016).

## Acknowledgements

Designed research SM, ARB, WAT; Performed research: SM, ARB, ETD, EAL, KDA, ADD, WAT; Contributed unpublished reagents/analytic tools: ETD, WAT; Analyzed data: SM, ARB, ETD, WAT; Wrote the paper SM, ARB, WAT.

The research reported in this manuscript was supported by the National Institute of Mental Health of the National Institute of Health under award numbers R01MH106568 and R01MH122843 to W.A.T. The content is solely the responsibility of the authors and the views expressed in this journal do not necessarily reflect the views or opinion of Catalent Pharma Solutions or the National Institutes of Health. The authors declare no competing financial interests.

